# Atomic-scale Characterization of Mature HIV-1 Capsid Stabilization by Inositol Hexakisphosphate (IP_6_)

**DOI:** 10.1101/2020.05.01.072686

**Authors:** Alvin Yu, Elizabeth M.Y. Lee, Jaehyeok Jin, Gregory A. Voth

## Abstract

Inositol hexakisphosphates (IP_6_) are cellular cofactors that promote the assembly of mature capsids of the human immunodeficiency virus (HIV). These negatively charged molecules coordinate an electropositive ring of arginines at the center of pores distributed throughout the capsid surface. Kinetic studies indicate that the binding of IP_6_ increases the stable life times of the capsid by several orders of magnitude from minutes to hours. Using all-atom molecular dynamics simulations, we uncover the mechanisms that underlie the unusually high stability of mature capsids in complex with IP_6_. We find that capsid hexamers and pentamers have differential binding modes for IP_6_. Ligand density calculations show three sites of interaction with IP_6_ including at a known capsid-inhibitor binding pocket. Free energy calculations demonstrate that IP_6_ preferentially stabilizes pentamers over hexamers to enhance fullerene modes of assembly. These results elucidate the molecular role of IP_6_ in stabilizing and assembling the retroviral capsid.

## Introduction

The capsids of the human immunodeficiency virus type-1 (HIV-1) encase and protect the retroviral genome from natural host defense mechanisms, but are transiently stable complexes. These capsids are assembled inside viral particles after proteolytic cleavage of the Gag polyprotein in a large-scale morphological transition of the virus termed maturation^1,2^ and must be disassembled prior to nuclear import of the virus’ genetic information into the host cell^3^. Structurally, immature Gag lattice precursors form incomplete spherical assemblies, composed primarily of hexamers of uncleaved Gag polyprotein^4,5^, while mature HIV-1 capsids are fullerene-like structures, composed of hexamer and pentamer building blocks of a single capsid domain (CA) protein^6,7^. There are exactly 12 pentamers in a single mature capsid that is topologically required to enclose the shell and impart curvature onto the CA lattice. Biochemical reconstitutions of CA assemblies show that CA can self-assemble into a wide range of morphologies, including helical tubes and fullerene cones^8^. CA lattices comprised solely of hexamers, however, form helical tubes, rather than fullerene structures.

Characterization of capsids with fullerene geometries^9^ that resemble those imaged by electron cryo-tomography within intact virions has been notoriously difficult, in part because of the short lifetimes under which capsids remain stable in isolation and the structural heterogeneity of the capsids themselves. Recent x-ray crystal structures demonstrate that pH-gated pores found at the center of hexamer subunits are distributed throughout the capsid surface, and contain an electropositive ring of arginines (R18) at the narrowest region of the pore^10,11^. Polyanions, including nucleotides, readily bind to the arginine ring. In particular, inositol hexakisphosphate (IP_6_) is a cellular polyanion that is abundantly present in the cytosol and enriched in packaged viral particles that bud off from the cell^12^. The addition of IP_6_ to *in vitro* reconstitutions of the capsid, drastically increases the stable lifetimes of the HIV-1 capsid from several minutes to over ten hours and promotes the assembly of fullerene capsids^12–14^. Viruses that fail to package IP_6_ do not mature properly and have unstable capsids^15^. Prior molecular simulations examined the conformational fluctuations of portions of the immature Gag lattice in the presence and absence of an IP_6_ ligand and found noticeable distortions of the six-helix bundle in the Gag spacer peptide 1 without IP_6_^13^. Mutagenesis studies have revealed that removal of electrostatic repulsion between CA domains at the arginine ring (R18A) causes assembly behavior consistent with an increased frequency of pentamer incorporation^8,16^. Despite these advances, there are no studies on whether IP_6_ interacts with pentamers, and the molecular mechanisms that underlie the increased stability and enhanced assembly of the mature capsid by IP_6_ remain unclear.

To investigate this, we performed long timescale all-atom (AA) molecular dynamics (MD) simulations of CA proteins with IP_6_ in bulk solution and analyzed the mechanisms by which IP_6_ binds to capsid pores. This analysis reveals that CA hexamers and pentamers have surprisingly different binding modes for IP_6_. To identify the protein-ligand interactions made during the IP_6_ binding process, we calculated three-dimensional density maps for IP_6_ that specify the regions the ligand occupies during dynamical simulation. These density maps show two unexpected, weaker-affinity sites for IP_6_ and CA. Free energy calculations of the binding process indicate that IP_6_ stabilizes pentameric CA states to a greater extent than hexameric CA states, and suggest IP_6_ binding shifts assembly conditions towards an increased population of pentamers to stabilize the fullerene capsid.

## Results

### CA hexamers and pentamers have differential IP_6_ binding modes

Molecular dynamics simulations of the complete mature HIV-1 capsid (∼100 million atoms, Figure 1A) are computationally costly. Instead, to investigate how IP_6_ stabilizes the capsid, we simulated individual hexamer and pentamer components of the capsid using unbiased, microsecond timescale AA MD simulations on the special-purpose computer hardware, Anton 2, at the Pittsburgh Supercomputing Center^17^. Similar to prior studies, IP_6_ was initially added to the bulk solvent, approximately 10 Å away from any non-water molecules^18,19^. Four systems each for the CA hexamer and pentamer were prepared with IP_6_ in randomized positions in bulk solvent, and selected for longer timescale simulations on Anton 2 (see: Materials and Methods for a complete description). In aggregate, the AA MD simulations totaled 32 µs for the hexamer and pentamer systems. In all cases, IP_6_ was found to bind to the pore of the capsomere.

**Figure 1.**
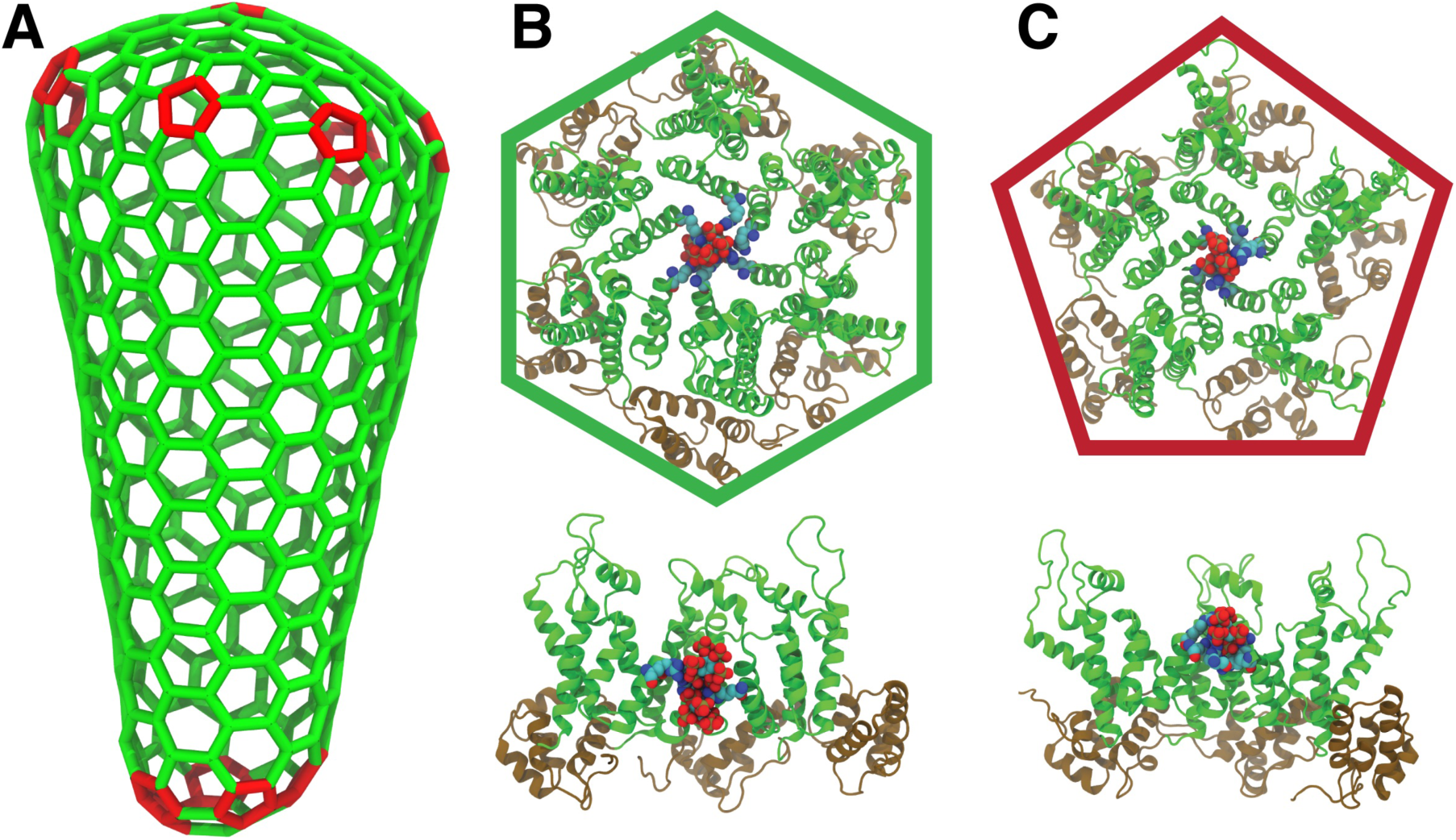
IP_6_ binds to the CA hexamer and pentamer. (**A**) The fullerene structure of HIV capsids contains exactly 12 pentameric defects. The geometry for the HIV-1 capsid was derived from cryo-electron tomography (cryo-ET) images of intact virions ^9^. Pentameric defects are colored red. (**B**) Two IP_6_ molecules bind to the R18 ring in the central pore of the CA hexamer after 0.73 µs of all-atom MD simulations. The bound complex is shown in a top-down and side view. (**C**) A single IP_6_ molecule binds to the R18 ring at a pore in the CA pentamer after 1.98 µs of AA MD simulations. The bound complex is shown in a top-down and side view. In the MD simulations for (**B**) and (**C**), IP_6_ molecules were initially placed in bulk solvent, ∼10 Å away from the CA domains. The NTD of CA is colored in green and the CTD of CA is colored in brown. R18 residues are shown in cyan.

After exhaustive MD simulation (∼ 4 µs per trajectory) of the CA hexamers, two IP_6_ molecules bind to a central arginine (R18) ring that lines the hexameric pore. A single IP_6_ ligand occupies the volume ∼ 3 Å above the R18 ring, with phosphate groups coordinated by the functional groups of the R18 sidechains, whereas a second IP_6_ molecule binds below the arginine ring, also coordinated by the R18 guanidino groups, proximal to the CA CTD (Figure 1B). These binding poses for IP_6_ are consistent with x-ray crystal structures of the mature CA hexamer in complex with IP_6_, which reveal electron densities for positions above and below the R18 ring^12,13^. Once bound, IP_6_ did not unbind from the R18 ring in the course of our simulations.

To our surprise, for all pentamer trajectories, only a single IP_6_ molecule bound to the R18 ring (Figure 1C). The closely packed positive charges of the R18 ring are mitigated by contacts with negatively charged phosphate groups on IP_6_. IP_6_ binds above the ring and does not interact significantly with protein residues below the R18 ring. Despite the overall structural similarities between capsomeres of the capsid shell, our simulations indicate that IP_6_ associates with hexamers and pentamers using distinctly different binding modes.

### Molecular mechanism of IP_6_ binding to CA hexamers and pentamers

The binding of IP_6,_ not only enables the *in vitro* assembly of fullerene capsids from concentrated CA solutions ([CA] > 1 mM), but also dramatically increases the lifetime of mature retroviral capsids synthesized in the laboratory from minutes to hours^20^. In the simulations, IP_6_ exits bulk solvent and can enter the constricted pore from either the N-terminal domain (NTD) or C-terminal domain (CTD) regions of the CA pentamer or hexamer. Snapshots of the binding process taken at various stages of ligand association are shown in Figures 2 and 3, and the full process is shown in Supplementary Movies 1 and 2.

**Figure 2.**
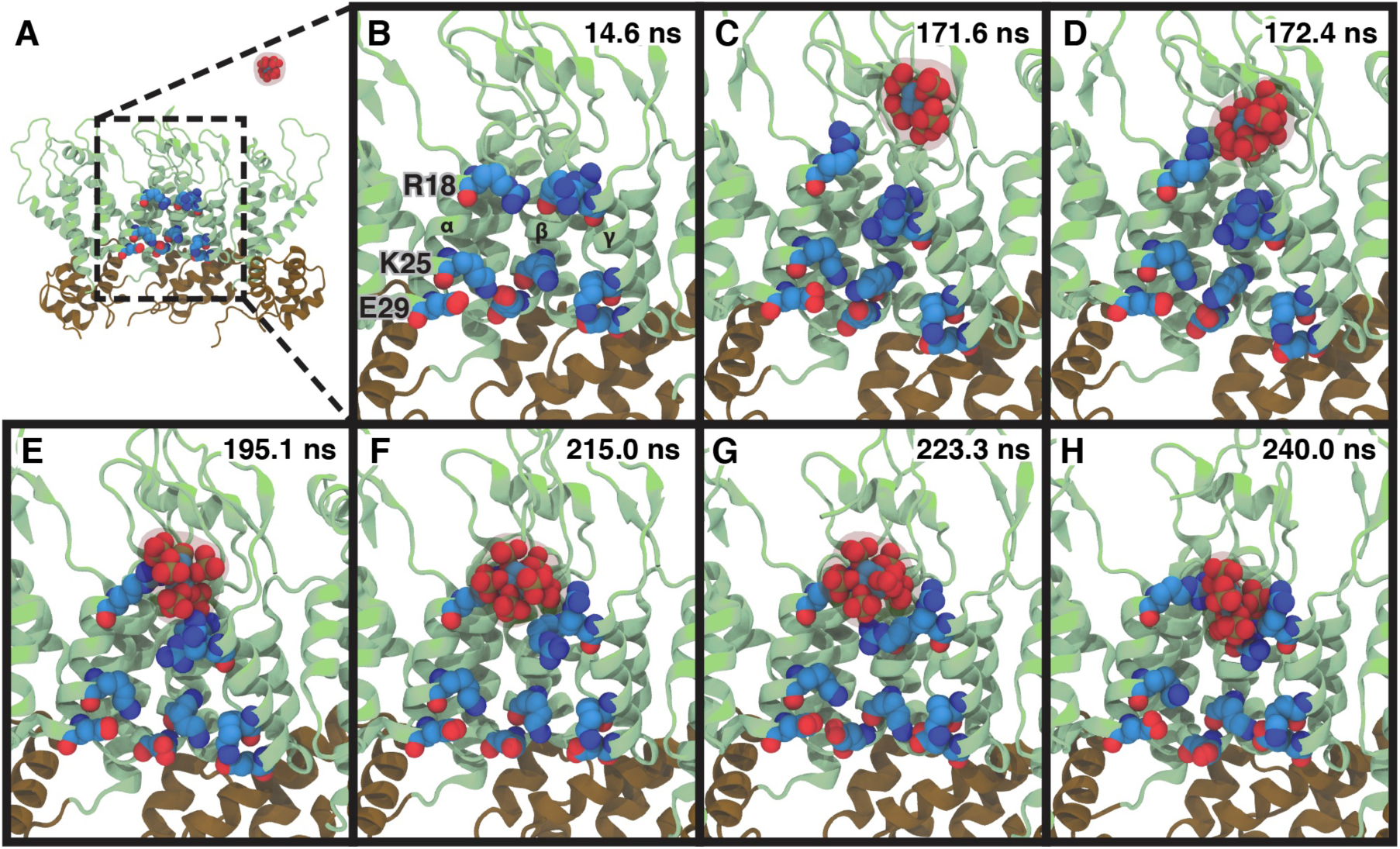
Mechanism of IP_6_ binding to CA pentamers. A series of snapshots from part of a MD trajectory shows the chemical interactions involved in IP_6_ binding to the CA pentamer. Three CA domains are shown in a side view of the pentamer, with helix H1 of the CA domain labeled as α, β, and γ, respectively. Two additional CA domains (δ and ε) are not shown for clarity. The R18 ring and residues (K25–E29) that form a salt bridge interaction proximal to the CTD are labeled. Time points are labeled in the upper-right corner of each panel. (**A**) The IP_6_ ligand is initially in bulk solvent. (**B**) Close-up view of the pore region in (**A**). (**C**) IP_6_ enters the pore past the beta-hairpin of CA, and an arginine side chain flips away from the R18 ring. (**D**) The R18 side chain coordinates IP_6_. (**E**) IP_6_ is coordinated to multiple R18 side chains. (**F**) IP_6_ shifts toward the R18 ring. (**G**) IP_6_ reorients in the binding pocket. (**H**) IP_6_ is bound to the central arginine ring with R18 side chains contacting the negatively charged phosphates.

**Figure 3.**
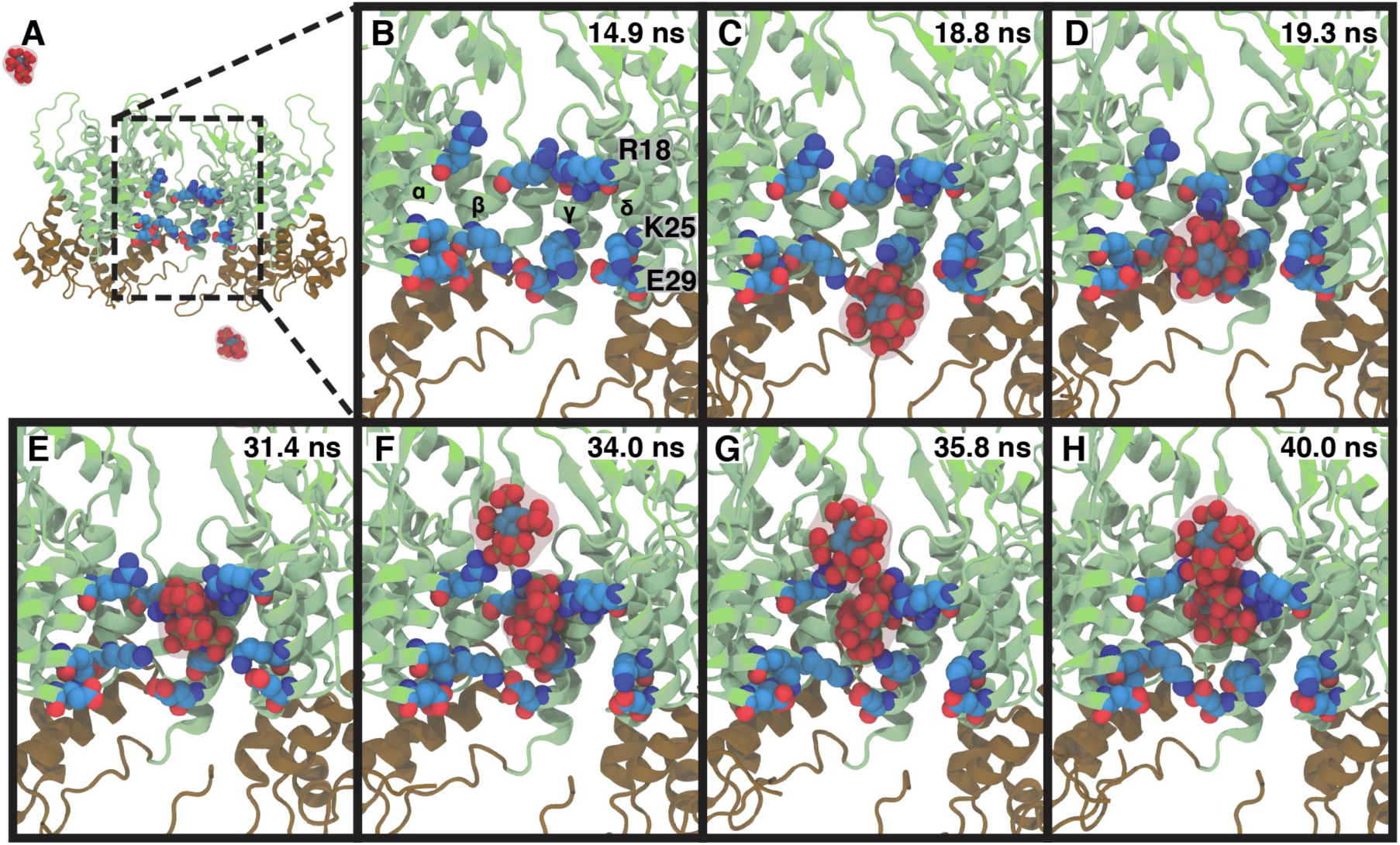
Mechanism of IP_6_ binding to CA hexamers. A series of snapshots from part of a MD trajectory shows the chemical interactions involved in IP_6_ binding to the CA hexamer. Four CA domains are shown in a side view of the hexamer, with helix H1 of the CA domain labeled as α, β, γ, and δ respectively. Two additional CA domains (ε and ζ) are not shown for clarity. The R18 ring and residues (K25–E29) that form a salt bridge interaction proximal to the CTD are labeled. Time points are labeled in the upper-right corner of each panel. (**A**) IP_6_ molecules are initially in bulk solvent (two are shown). (**B**) Closed-up view of the pore region in (**A**). (**C**) An IP_6_ ligand enters the pore region from the CTD of CA. The K25–E29 salt bridge interaction breaks, and K25 contacts IP_6_. (**D**) A R18 sidechain flips towards the CTD to coordinate IP_6_. (**E**) IP_6_ binds at the CTD side of the R18 ring. (**F**) A second IP_6_ ligand enters the pore region, coordinated by a R18 side chain. (**G**) Multiple R18 side chains contact the second IP_6_ ligand. (**H**) Two IP_6_ molecules are bound to the R18 ring. Contacts between IP_6_ and K25 are broken, and salt bridge interactions between K25 and E29 reform.

For the CA pentamer, IP_6_ initially diffuses past the beta-hairpin loop in the NTD. The sidechains of the R18 ring are typically modeled to be planar in x-ray crystal structures and cryo-EM maps with torsional angles of (χ_1_ = −74°, χ_2_ = −156°, χ_3_ = 71°, χ_4_ = 84°) (Figure 2A–B), yet prior to binding, the pentameric complex, surprisingly, shifts asymmetrically, involving a vertical separation of ∼8 Å between the Cα carbons of R18 on helix H1 of the α and β domain, and the CA domains twist helically upwards with β < γ < δ < ε < α along the axis of the pore (Figure 2B– C). The pore arginine of the α-CA domain flips away from the plane with rotations about the R18 side chains (χ_1_ = −70°, χ_2_ = 57°, χ_3_ = 60°, χ_4_ = −179°), likely reflecting unfavorable electrostatic interactions between the guanidino groups (Figure 2C). The arginine side chain coordinates a phosphate group on IP_6_ (Figure 2D), as IP_6_ binds to the pocket above the R18 ring. Additional arginine groups from the remaining CA domains contact the uncoordinated phosphate groups of the ligand, as IP_6_ binds into the central pore, and the R18 ring relaxes to a planar configuration (Figure 2E–H).

In the CA hexamer, the overall complex does not twist asymmetrically, and IP_6_ enters the pore initially from the CTD (Figure 3A–C). Salt-bridge interactions between K25–E29 at the base of helix H1 break, and the lysine side chains interact with IP_6_ (Figure 3C). One of the pore arginines of the β-CA domain rotates downward to contact a phosphate group on IP_6_ and forms a metastable complex between the protein and ligand, in which two IP6 phosphate groups are coordinated at both ends by R18 and K25 (Figure 3D–E). A second IP_6_ molecule enters the pore from the NTD region, anchored by an uncoordinated arginine in the R18 ring (Figure 3F). Finally, interactions between the first IP_6_ molecule and K25 break, as both IP_6_ ligands relax into the central binding sites above and below the R18 ring (Figure 3G–H). X-ray structures show an iris-like opening of the CA hexamer pore in response to acidic pH conditions, hypothesized to occur by protonation of a critical histidine residue (H12), and a conformational change in the β-hairpin that gates the pore^10^. In the simulations however, IP_6_ bound in the same fashion to the R18 ring for CA complexes containing both protonated and de-protonated H12, and either in the open or closed conformations. The β-hairpin regions were quite flexible, suggesting that opening or closing of the proposed gate is driven more by small changes in the conformational ensembles adopted by the β-hairpin rather than abrupt structural changes.

### Ligand density maps reveal alternate binding pockets

Transient metastable interactions between the protein and ligand can contribute substantially to the association process by lowering the energetics of intermediate states along the binding pathway. Prior simulations of other systems have discovered a wide range of dynamical mechanisms utilized for binding, which do not involve direct interactions between the ligand and the binding site, including protein conformational changes that open up previously inaccessible binding pockets^21–23^, the flipping of ligand configurations between several bound poses^18,24^, and metastable electrostatic interactions that guide ligands into the binding pocket^18,19,25,26^. To assess the protein-ligand interactions that form during our simulations, we computed the spatial distribution occupied by the ligand’s non-hydrogen atoms, or in other words a three-dimensional map of the ligand density (Figure 4).

**Figure 4.**
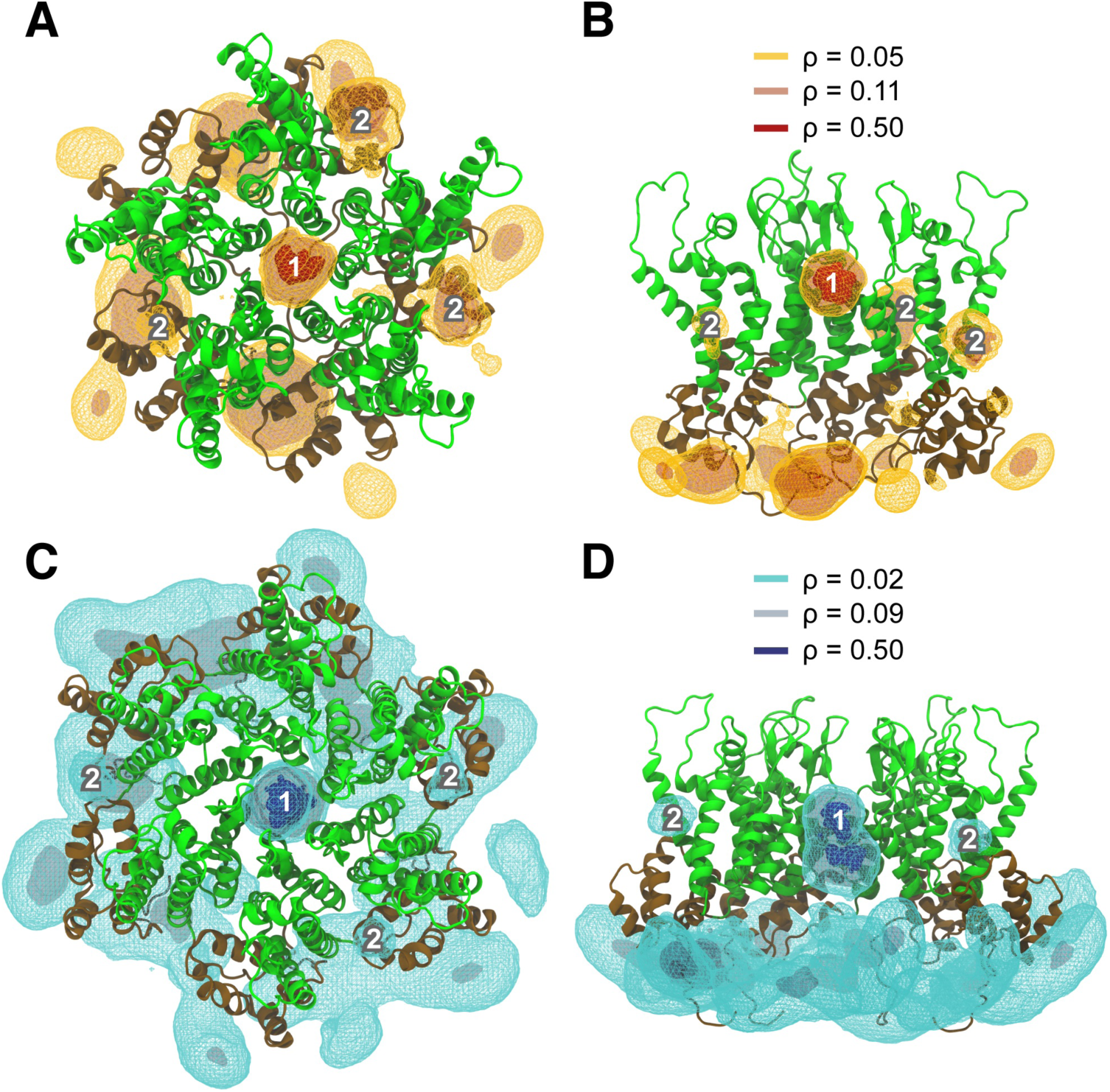
3D ligand density maps for inositol hexakisphosphate. The ligand densities are contoured at *ρ* = 0.05, 0.11, and 0.50 from orange to red, overlaid onto the structure of the CA pentamer, in a top-down view (**A**) and side view (**B**). Ligand densities are contoured at *ρ* = 0.02, 0.09, and 0.50 from light blue to dark blue, overlaid onto the structure of the CA hexamer in a top-down view (**C**) and side view (**D**). Densities contributing to interactions at sites 1 and 2 are labeled in the figure, whereas densities proximal to the CA CTD (brown) form a third interaction site.

The ligand density maps reveal three sites of interaction between IP_6_ and the CA domain pentamer (Figure 4A and 4B). The primary binding site (Site 1) has the highest density for the ligand and is positioned at the central pore of the pentamer, stabilized by interactions between IP_6_ and the R18 ring. Intriguingly, the simulations also showed a second, lower-affinity binding site (Site 2) at an intermolecular interface between the NTD of one CA domain, and the CTD of the adjacent CA domain. This secondary site is the binding pocket for several drug molecules that target the HIV capsid and inhibit viral replication including BI-2^27^, PF74^28^, and GS-CA1^29^. Although the precise mechanisms of action for these compounds are currently unclear, several studies have suggested that small-molecule binding at this site may overly stabilize the capsid and perturb normal capsid disassembly processes, interfere with CA domain assembly processes that form intact capsids, or compete with nuclear import factor binding to the capsid, which is required for integration of the viral genome into the host cell^30,31^.

Significant density is also observed at a third site (Site 3) proximal to the CTD and consists of interactions between the flexible CA CTD-tail and the negatively charged IP_6_ ligand. There are no known drug molecules that currently bind to this site, which is a cleavage by-product of the retroviral aspartyl protease (e.g., HIV protease) during virion maturation. The aspartic acid moieties in the catalytic site of HIV protease may in part explain the affinity of the CTD tail for negatively-charged molecules; although an alternative hypothesis is that the electrostatic properties of the CTD-tail allow the CA CTD to form interactions with viral RNA that are necessary for recruiting and containing RNA in the mature capsid.

Ligand density maps for the CA domain hexamer are largely similar to that of the pentamer, containing the same sites and interactions with the IP_6_ ligand (Figure 4C and 4D). One noticeable difference, however, is that the primary site (Site 1) in the hexamer is split into two continuous bodies corresponding to binding sites for IP_6_ above and below the R18 ring at high contour levels for the density (ρ = 0.5), whereas the map in the pentamer contains only a single density for Site 1 above the R18 ring. This is consistent with our prior result that hexamers can bind two IP_6_ molecules, whereas pentamers bind only a single IP_6_ molecule. Our ligand densities for Sites 2 and 3 are not entirely either five-fold or six-fold symmetric around the pentamer or hexamer respectively, owing to challenges in converging sampling statistics for IP_6_ and CA interactions at each of the binding sites. The binding sites discovered from our density analysis, however, agree well with known interactions at the CA domain, such as interactions with the capsid inhibitors (i.e. BI-2, PF74, and GS-CA1) and HIV protease. Our AA MD simulations did not contain prior information on these interactions, highlighting the potential of such simulations to detect novel binding sites for other molecules.

### Free energy landscapes for IP_6_ binding

To quantify the energetics of the binding process, we computed the free energy landscape, i.e., the potential of mean force (PMF), for IP_6_ binding to CA hexamers and pentamers using umbrella sampling simulations, in which a one-dimensional order parameter was used to describe the transport of IP_6_ to the R18 ring. The 1D order parameter, ξ, is defined as the projection of the displacement between the center-of-mass of the IP_6_ ligand and the center-of-mass of the backbone atoms of the R18 ring onto the vertical pore axis, and is shown for the CA hexamer and CA pentamer in Figure 5A and 5B, respectively.

**Figure 5.**
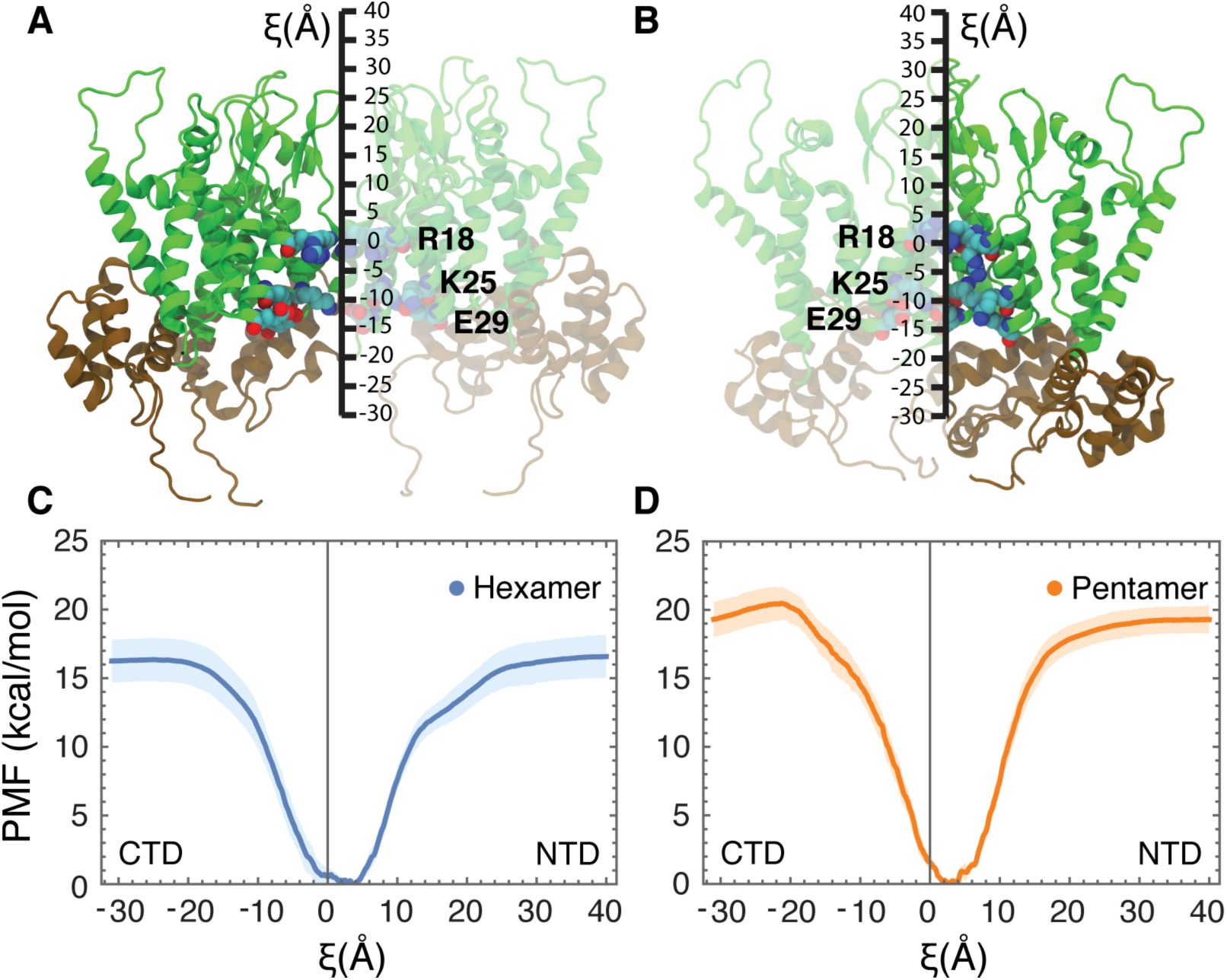
The potential of mean force (PMF) for IP_6_ binding. The order parameter, ξ, was used to describe IP_6_ transport to the R18 ring. ξ is defined as the projection of the center-of-mass displacement between IP_6_ and R18 residues onto the pore axis, and is shown schematically for the CA hexamer (**A**) and the CA pentamer (**B**). The NTD is colored in green, whereas the CTD is colored in brown. Residues R18, K25, and K29 are labeled. (**C**) The 1D PMF for the CA hexamer is shown in blue. The shaded light-blue regions denote the error in the PMF determined by block-averaging. (**D**) The 1D PMF for the CA pentamer is plotted in orange. The shaded light-orange regions denote the error in the PMF determined by block-averaging. The pore axis is oriented such that positive values correspond to the CA NTD, whereas negative values correspond to the CA CTD.

Both PMFs (Figure 5C and 5D) show that IP_6_ binding alleviates significant electrostatic repulsion between the R18 sidechains and contain deep free energy minima proximal to the R18 ring (*ξ* = 0 Å). The hexamer PMF has a broad free energy basin reflecting a larger pore volume for IP_6_ diffusion, and a global free energy minimum positioned at *ξ* = 3.*7* Å. There is a second, local minimum in the hexamer PMF at *ξ* = −0.8 Å corresponding to the binding site below the R18 ring, which is slightly less stable relative to the global free energy minimum (Δ*G* = 0.6 kcal/mol). This is consistent with electron density maps derived from x-ray crystal structures of CA hexamers in complex with IP_6_ that revealed stronger densities for binding sites above the ring than below the ring^12,13^. In contrast, the pentamer PMF has a narrower free energy basin owing to a more restricted and compact helix H1 region below the R18 ring in the CA pentamer as compared to that of the CA hexamer and has a single overall free energy minimum located at *ξ* = *2*.*7* Å. Ligand entry from the CTD side of the pentamer is unfavorable compared to entry from the NTD region, and conformational states in which IP_6_ is below the R18 ring (*ξ* < 0 Å) are destabilized (Δ*G* > 1.6 kcal/mol). In both hexamer and pentamer PMFs, the CA domain β-hairpin (*ξ* ∼ 5– 15 Å) does not form significant free energy barriers to ligand entry.

As the ligand exits into bulk solvent (*ξ* > 25 Å and *ξ* < −20 Å), both PMFs plateau to a constant value as expected, since the protein and ligand no longer interact. The total free energy difference of transporting IP_6_ to the R18 ring is greater in the pentamer (Δ*G* = 19.3 kcal/mol) than in the hexamer (Δ*G* = 16.4 kcal/mol), indicating that IP_6_ binding stabilizes CA pentamers more than CA hexamers in the fullerene shell. The calculated binding free energies are consistent with that of 4–6 Arg–phosphate interactions, which have higher stabilities^32–34^ than that of Arg–Glu interactions (Δ*G*_RE_ = ∼4– 5 kcal/mol for a single pair^35,36^). Hexamer configurations can form an additional Arg–phosphate interaction compared to pentamer configurations; however the smaller pentamer pore facilitates closer coordination of IP_6_ phosphate groups by R18 side chains. Negative-stained electron microscopy studies demonstrated that removal of the charged arginine ring by an R18A mutation forms CA assemblies with spherical, fullerene, and spiral structures, hypothesized to occur from an increased frequency of pentamer incorporation into the CA hexagonal lattice^8,37^. Overall, our findings suggest that IP_6_ preferentially binds to and stabilizes CA pentamers over CA hexamers as a result of the more compact and closely packed H1 pore helices in the pentamer and a subsequent greater electrostatic repulsion at the R18 ring. Given the higher number of hexamers in the capsid lattice, a large amount of IP_6_ in total should be found among the hexamers.

## Discussion

Collectively, our results suggest that CA hexamer and pentamer building blocks of the fullerene capsid bind IP_6_ with rather different binding modes. At high IP_6_ concentrations (i.e., [IP_6_]:[CA] = 1:3), hexamers can bind IP_6_ at a secondary site below the R18 ring, whereas at low IP_6_ concentrations ([IP_6_]:[CA] = 1:6), IP_6_ alternates between the two binding sites; although the site above the R18 ring is lower in free energy (Δ*G* = −0.6 kcal/mol). Experimental estimates using radiolabeled IP_6_ ligands suggest that ∼300 IP_6_ molecules are packaged per viral particle, which contains ∼1,500 CA domains, indicating that the latter case is more likely to occur^10^. The molecular pathways by which IP_6_ binds involves a conformational twist in the pentamers, which may also be aided by the inclusion of pentamers at highly curved regions of the capsid surface. Binding below the R18 ring in pentamers is less favorable since pentamers have a compact and tighter pore relative to that of hexamers. Ligand density calculations show that interactions at the R18 ring are consistent with the differential binding modes and reveal two additional sites of IP_6_ interaction, one of which is a previously known capsid-inhibitor binding pocket and the other consists of a novel interaction between the CTD tail and negatively charged molecules that could reflect capsid and RNA interactions.

The computed PMFs demonstrate that IP_6_ binding stabilizes individual pentamers more than individual hexamers. Since CA spontaneously assembles into both helical tubes and fullerene cones *in vitro* ^38^, and pentameric defects are required to produce fullerene cones but are not present in helical tubes, this result indicates the binding of IP_6_ shifts the equilibrium population of hexamers and pentamers towards increased pentamer formation, thereby promoting the fullerene mode of assembly. It is of note to recognize that the pathways discussed characterize the binding of IP_6_ to preformed mature capsid complexes and not the molecular pathways for the co-assembly of CA and IP_6_, which may differ. Further simulations that target the co-assembly of CA and IP_6_ will likely require coarse-grained molecular simulation techniques to overcome the long timescale barriers associated with biomolecular assembly phenomena^39–41^. The computed thermodynamic quantities however are state functions that do not change in response to the particular path taken, so the IP_6_ interactions revealed by the ligand density maps are therefore general.

Since inositol phosphates plays important physiological roles in promoting and stabilizing the assembly of the HIV-1 capsid, could the removal of IP_6_ be involved in disassembly processes? One possible mechanism is that acidic conditions simultaneously trigger protonation of IP_6_ phosphate groups and H12, thus reducing the affinity of IP_6_ for the R18 ring and opening of the capsid pores. Although pH has been known to trigger the release of genetic material in viruses by acidification of endosomal vesicles during intracellular transport^42^, it is unclear whether HIV capsids are exposed to similarly low pH conditions during infection in either endocytic or membrane fusion pathways^43^.

In this work, we have elucidated the molecular mechanisms responsible for the stabilization of mature capsids by inositol hexakisphosphate through extensive AA MD simulations. Additional investigations can be carried out to further probe how retroviruses exploit environmental cues to initiate replication processes.

## Materials and Methods

### All-atom models for the CA Hexamer, Pentamer, and IP_6_

The initial atomic models for the CA hexamer and pentamer were constructed from the x-ray crystal structures for the CA hexamer in the closed and open conformations (PDB ID: 3H47, 5HGL, respectively), and the cryo-electron microscopy (cryo-EM) structure for the CA pentamer in the closed conformation (PDB ID: 5MCY). A homology model for the open CA pentamer was constructed based by aligning individual CA domains and transplanting the beta-hairpin gate (resid 1–16) from the open CA hexamer onto the closed pentamer. Missing amino acid residue backbones were built using MODELLER^44^, and missing side chains were built using SCWRL4^45^. The hexamer and pentamer systems, which contained a total of 287,048 and 286,715 atoms respectively, were solvated and neutralized by adding Na^+^ and Cl^-^ ions to the bulk solution until the salt concentration was 150 mM NaCl. Periodic boundary conditions were imposed on an orthorhombic unit cell of approximately 141 Å × 141 Å × 141 Å for both the hexameric and pentameric systems. The all-atom potential energy function CHARMM36m^46,47^ for proteins and the TIP3P ^48^ potential energy function for water were used. Electrostatic interactions were computed using the particle mesh Ewald (PME) algorithm and short-range, non-bonded interactions were truncated to 12 Å. Harmonic restraints were applied to the center-of-masses of two groups of residues (resid 179–189 and resid 200–207) in α-helices, H9 and H10, at the CTD interface between distinct hexamer and pentamer subunits to maintain the relative position of the CTD interface derived from an atomic model of the complete HIV-1 capsid^9^. For these restraints, a light force constant of *k* = 0.5 kcal/mol was used. The solvated system was energy minimized and equilibrated under constant pressure and temperature (NPT) conditions at 1 atm and 310 K with a 2-fs timestep. Minimization and equilibration procedures were performed using the AA MD simulation package NAMD 2.13^49^.

### MD Simulations for IP_6_ Binding

IP_6_ molecules were initially placed at random positions and orientations in bulk solvent, approximately 10 Å away from any non-water molecules. 15 IP_6_ molecules were used to increase sampling times. The IP_6_ was parameterized using the CHARMM General Force Field (CGenFF)^50^. For each CA domain system, the histidine residues, H12, were completely protonated (HSP) or de-protonated (HSD). Salt concentration was adjusted for the ligand charge to maintain 150 mM NaCl and an electrostatically neutral system. After a 50 ns equilibration run, eight systems consisting of hexamer/pentamer, open/closed, and H12 protonated/de-protonated combinations were prepared for simulation on Anton 2 at the Pittsburgh Supercomputing Center^17^. For all production simulations, bond lengths for hydrogen atoms were constrained using the M-SHAKE algorithm ^51^. An r-RESPA integrator was used with a timestep of 2 fs^52^; long-range electrostatics were computed every 6 fs. Long-range electrostatics were calculated using the k-space Gaussian split Ewald method^53^. Short-range interactions including van der Waals and short-range electrostatics were truncated at 9 Å. Simulations in the constant NPT ensemble were performed using a chained Nosé-Hoover thermostat at 310K and isotropic Martyna-Tobias-Klein barostat at 1 atm^54^. Each trajectory (4 CA hexamer and 4 CA pentamer) was simulated until no additional binding events were observed for 2 µs. In aggregate, simulation time was 32 µs for the CA hexamer and pentamer systems.

### Ligand density calculations

The trajectories from production-level simulations were subsampled at 0.12 ns intervals and aggregated into pentamer or hexamer systems. For each frame, the atomic system was aligned by minimizing the RMSD of the protein backbone atoms with respect to the crystallographic structure. Cartesian coordinates for the non-hydrogen atoms of IP_6_ were recorded, and the density of atomic positions 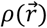 was calculated, using a hard-sphere van der Walls approximation on a discretized grid with a spacing of 0.5 Å × 0.5 Å × 0.5 Å. The 3D ligand density map was subsequently contoured at *ρ* = 0.05, 0.11, 0.50 for the CA pentamer systems and *ρ* = 0.02, 0.09, 0.05 for the CA hexamer systems.

### Umbrella Sampling

The free energy landscape (PMF) of IP_6_ binding was computed using an umbrella sampling strategy^55^. All systems contained a single IP_6_ molecule and either a CA hexamer or pentamer in the open conformation. A 1D order parameter, ξ, is used to describe the relative distance to the R18 ring. ξ specifies the projection of the displacement vector between the center-of-mass of the IP_6_ ligand 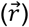 and center-of-mass of the backbone atoms of the R18 ring 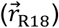 onto the z-axis (e.g. 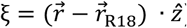). The umbrella sampling windows consisted of CA domain complexes and the IP_6_ ligand positioned in 0.2 Å increments along ξ. The initial coordinates were obtained using targeted MD simulations in NAMD, using the solvated CA hexamer and pentamer systems in the open conformation with a protonated H12 residue. Three hundred sixty umbrella sampling windows were used to compute the PMFs. Harmonic biasing potentials with a force constant of 20 kcal/mol/Å^2^ centered on ξ were used. All other restraints and interactions were computed as described during the equilibration phase. The total simulation time for the CA hexamer is 28.8 µs. The total simulation for the CA pentamer is 28.8 µs. Each PMF, *W*(ξ), was computed using the weighted histogram analysis method (WHAM) to unbias and recombine the sampled distribution functions from all windows^56,57^. The statistical uncertainty in each PMF was evaluated using the approach of block averaging^58^. For each PMF, the time series was divided into 5 blocks, and WHAM was used to calculate a PMF from the data in each block. The standard deviation of the 5 PMFs was reported. Using 5–15 blocks all gave qualitatively similar results.

## Author Contributions

A.Y. and G.A.V designed research. A.Y. performed research. A.Y., E.M.Y.L. contributed new reagents or analytic tools. A.Y., E.M.Y.L., J.J., G.A.V. analyzed data. A.Y., E.M.Y.L., J.J., and G.A.V wrote the paper.

## Acknowledgements

This work was supported by the National Institute of Health grant P50 AI150464 for the Center for the Structural Biology of Cellular Host Elements in Egress, Trafficking, and Assembly of HIV. Computational resources were provided by Frontera at the Texas Advanced Computer Center funded by the National Science Foundation grant (OAC-1818253). Anton 2 computer time was provided by the Pittsburgh Supercomputing Center (PSC) through grant R01 GM116961 from the National Institute of Health. The Anton 2 machine at PSC was generously made available by D.E. Shaw Research. A.Y. gratefully acknowledges support from the National Institute of Allergy and Infectious Diseases of the National Institute of Health under grant F32 AI150208.

